# Rapid Cas13a-based *pen*A genotyping for cefixime susceptibility in *Neisseria gonorrhoeae*

**DOI:** 10.64898/2026.02.26.708400

**Authors:** Thi Hai Yen Nguyen, Sakshi Garg, Gordon Adams, Sreekar Mantena, Nisha Gopal, Ho-Jun Suk, Jeffrey D. Kalusner, Pardis C. Sabeti, Jacob E. Lemieux, Lao-Tzu Allan-Blitz

## Abstract

**Background:** Antimicrobial resistance in *Neisseria gonorrhoeae* is an urgent public health threat. Resistance-guided therapy can assure appropriate treatment and reintroduce alternative therapeutic options by identifying genetic predictors of resistance. Mosaicism at codons 375-377 of the *pen*A gene are associated with cefixime resistance. Rapid, field-deployable assays for predicting cefixime susceptibility are lacking.

**Methods:** We used a machine-learning algorithm to develop a CRISPR Cas13a-based assay to detect the absence of mosaicism at codons 375-377 of the *pen*A gene combined with isothermal amplification in a single reaction. We integrated the assay onto a portable fluorescence-based platform. We evaluated performance using cultured isolates and compared results with PCR genotyping and phenotypic antimicrobial susceptibility testing. We also assessed feasibility of reagent lyophilization for cold-chain–independent deployment.

**Results:** Among 40 *N. gonorrhoeae* isolates, the Cas13a *pen*A assay demonstrated 100% concordance with PCR genotyping and 92·5% (95% CI 79.6-98.4%) concordance with phenotypic cefixime susceptibility. Median time to detection was 12 minutes (IQR 5 minutes). The lyophilized detection system detected all 12 isolates with a median time to detection of 45·0 minutes (IQR 40-45) compared to 45·0 minutes (IQR 35-50) for the positive aqueous control, although peak fluorescence was higher for the aqueous control (*p*<0.01).

**Conclusion:** The Cas13a assay was rapid and demonstrated strong correlation with genotypic and phenotypic cefixime susceptibility in *N. gonorrhoeae*, while a lyophilized assay retained functionality.

## Introduction

Antimicrobial resistance in *Neisseria (N*.*) gonorrhoeae* infection is an urgent public health threat.^1,2^ There were over 80 million new cases of gonorrhoeae worldwide in 2020,^3^ with the highest prevalence among low-resource settings.^4^ Further, *N. gonorrhoeae* has now developed resistance to all antimicrobials used in its treatment, including recent reports of high rates of resistance to ceftriaxone – the last line empiric therapy.^5-7^ Because routine culture is not feasible, and because results would not be available rapidly enough to inform care, the diagnostic paradigm relies on nucleic acid amplification testing, which provides no information on antibiotic susceptibility. Thus, all *N. gonorrhoeae* infections are treated with a single regimen, which applies selective pressure toward the emergence of resistance.^8^ Furthermore, low-resource settings lack laboratory infrastructure to support nucleic acid amplification testing. Instead, such areas utilize syndromic management, which misses the high proportion of asymptomatic cases and results in antibiotic overuse among symptomatic patients,^9^ further driving resistance.

Rapid prediction of antimicrobial susceptibility to guide care, known as resistance-guided therapy, is possible via molecular assays that detect the genetic determinants of resistance.^10^ The absence of mutation in codon 91 of the gyrase A (*gyr*A) gene was more than 98% sensitive and 98% specific for predicting ciprofloxacin susceptibility.^11^ *Gyr*A genotyping assays are increasingly available using polymerase chain reaction (PCR), recombinant polymerase amplification (RPA) and even CRISPR.^12-14^ Resistance-guided therapy is increasingly being used for other pathogens as well, including *Staphylococcus aureus* and *Helicobacter pylori*.^15,16^ The 2021 U.S. Centers for Disease Control and Prevention Sexually Transmitted Infection treatment guidelines permit the use of ciprofloxacin in settings where rapid genotyping assays are available.^17^ However, the effectiveness of resistance-guided therapy to mitigate the selective pressure towards ceftriaxone resistance is enhanced when molecular detection platforms incorporate genetic targets that predict resistance to additional antibiotics.^18^

Prior work has demonstrated that the absence of mutation at any of 6 loci in the *pen*A gene was between 95-99% sensitive for predicting susceptibility to cefixime, with the absence of mosaic insertions in codon 375-377 of particular importance.^19,20^ The World Health Organization lists cefixime as an acceptable alternative first-line agent for uncomplicated gonorrhea,^21^ thus rapid prediction of cefixime resistance may be able to avert treatment failures. We aimed to develop a low-cost, field-deployable system for rapidly determining the absence of *pen*A mosaicism in *N. gonorrhoeae*. To do so, we leveraged the Specific High-Sensitivity Enzymatic Reporter UnLOCKing (SHERLOCK) platform we previously utilized to develop assays for *N. gonorrhoeae* detection and *gyr*A genotype prediction.^13,22^ That system employs the isothermal amplification, T7-polymerase mediated transcription, and RNA-guided CRISPR enzyme Cas13a detection via cleavage of a quenched reporter.^23^ We further aimed to utilize a field-deployable device for simultaneous amplification, transportation and detection. In addition, we aimed to lyophilize or freeze-dry reagents to permit cold-chain-independent storage and facilitate deployment in field settings.

## Methods

### Cas13a Guide RNA and RPA Primer Designs

To design Cas13a guide RNA sequences, we used a machine-learning tool known as Building Artificial Diagnostic Guides by Exploring Regions of Sequences (BADGERS).^24^ BADGERS employs a predictive model of guide-target activity to explore a fitness landscape of candidate guide sequences and design those with optimal on-target and minimal off-target activity. As input, we provided 2,274 unique *N. gonorrhoeae* genomes (n=1,675 with mosaicism in the *pen*A gene) and specified codons 375-377 as the design window. We provided nucleotide sequences of *N. gonorrhoeae* isolates with mosaicism at codons 375-377 in the *pen*A gene as the first target set and nucleotide sequences of *N. gonorrhoeae* isolates without mosaicism at those positions as the second target set. We selected potential candidate CRISPR guide RNAs (gRNAs) for the absence of *pen*A mosaicism based on scores generated by BADGERS for predicted fitness as well as predicted mean on and off target binding affinity while also attempting to ensure heterogeneity in start codon position.

We then developed two forward and reverse RPA primer sets (P1 and P2) flanking the mosaic region using PrimerBlast (National Center for Biotechnology Information). We designed the primers to yield amplicons ranging from 140 to 200 base pairs in length, with melting temperature between 58°C and 68°C. To enable *in vitro* transcription, we appended a T7 RNA polymerase promoter sequence (5’-GAAATTAATACGACTCACTATAGG-3’) to the 5’ end of each forward primer. We evaluated each gRNA-primer pair in a one-pot SHERLOCK reaction (see below) to assess both amplification efficiency and discrimination between target and non-target sequences. We used the synthetic *pen*A DNA that included the primer binding regions and gRNA binding site.

### SHERLOCK One-Pot Reaction

We performed the one-pot SHERLOCK reactions under standard Streamlined Highlighting of Infections to Navigate Epidemics (SHINE) conditions as previously described.^25^ Briefly, the master mix contained 1 x SHINE buffer (20 mM HEPES pH 8·0 with 60 mM KCL and 5% polyethylene glycol (PEG), 45 mM *Lwa*Cas13a (Genscript, Z03486-100), 1 U/µL murine RNase inhibitor (New England Biolabs, USA), 10 U/µL T7 RNA polymerase (Lucigen Corporation, USA), 136 mM RNaseAlert substrate v2 (Thermo Fisher Scientific, USA), and 2 mM of each ribonucleotide (rNTP) (New England Biolabs, USA). Detailed information on reagents, including suppliers and stock concentrations, is provided in Supplemental Tables 1 and 2.

We used the prepared master mix to resuspend lyophilized TwistAmp Basic Kit RPA pellets (one pellet per 73·42 µl master mix volume), followed by magnesium acetate MgAOc (TwistDx, United Kingdom) to a final concentration of 14 mM, which served as the sole magnesium cofactor. The mixture also included assay-specific concentration 320 nM of each forward and reverse RPA primers and 22·5 nM gRNA. We added target DNA to the final master mix at a 1:4 master mix-to-sample ratio. For evaluating gRNA and primer set performance, we ran the assays in triplicate on the Cytation5 plate reader (Agilent Technologies, USA) at 37°C, measuring real-time fluorescence (excitation 485nm; emission 528 nm) every 5 minutes for 3 hours.

After selecting the gRNA and primer set, we evaluated performance of the assay on cultured isolates using the DxHub platform (DxLab Inc., United States; Manufactured under contract by Axxin, Australia). The DxHub is a portable, standalone platform that provides independent testing of up to eight individual tubes, with isothermal incubation between 38°-72°C and dual-channel fluorescence detection in real-time. We ran reactions for 60 minutes at 38°C and measured fluorescence kinetics every 20 seconds.

### Quantitative Polymerase Chain Reaction

We designed a quantitative polymerase chain reaction (qPCR) system for confirmatory genotyping of the mosaic *pen*A region codons 375-377. We designed the assay such that the forward primer overlapped the non-mosaic consensus sequence; thus, any amplification would indicate the absence of mosaicism. The forward and reverse primer sequences targeting codons 375-377 of the *N. gonorrhoeae pen*A gene were 5’-GCTGAATACGCAGCCTTATAAAATCGG-3’ and 5’-TTTCTCAACAAACCTGCAGTTTCCC-3’, respectively.

We used 1 x FastStart SYBR Green Master Mix (Sigma-Aldrich, USA), 0·5 µM of each primer, and DNA template in a 1:9 template-to-master mix ratio. We adjusted the final reaction volume to 10 µl with nuclease-free water and performed reactions in triplicate on a 384-well plate using a QuantStudio 6 system (Applied Biosystems, USA). We used the following thermal cycling conditions: initial denaturation at 95°C for 3 min, 40 cycles at 95°C for 15 sec, 60°C for 1 min, and 72°C for 1 min; followed by a final extension at 68°C for 2 min. We measured fluorescence signals during the extension phase.

### Cas13a-based penA genotyping on N. gonorrhoeae isolates

We evaluated the selected Cas13a assay performance on purified *N. gonorrhoeae* isolates. All isolates were stored at −80°C in Microbank vials (Pro-Lab Diagnostics, Canada), which contained cryoprotectant beads designed to preserve bacterial viability during long-term storage. We cultured *N. gonorrhoeae* isolates on Thayer-Martin agar at 37°C in 5% CO_2_ for 24-48 hours. All isolates had previously been phenotypically characterized by standard antimicrobial susceptibility testing methods. We extracted DNA from the isolates using the DNeasy Blood and Tissue Kit (Qiagen, Germany) in accordance with the manufacturer’s instructions.

We tested each isolate with the Cas13a *pen*A mosaic assay on the DxHub and qPCR. The DxHub has a programmable positivity threshold that can integrate fluorescence amplitude and the rate of rise of the fluorescence signal. As we had not yet established a positivity threshold for the new assay, we defined a positive fluorescence signal as a peak at least three standard deviations above the negative control. We defined the time at which peak fluorescence occurred as the point at which the slope of the fluorescence amplitude equaled zero. We defined the time to positivity as the time to ½ peak fluorescence (analogous to the maximal slope).

The Clinical and Laboratory Standards Institute defines cefixime phenotypic resistance as a minimum inhibitory concentration (MIC) of ≥ 0·25 µg/mL with intermediate susceptibility as an MIC of 0·125 µg/mL.^26^ We considered isolates with an MIC ≥ 0·125 µg/mL as non-susceptible for comparison purposes, and compared the performance of Cas13a *pen*A mosaicism determination with 1) qPCR results and 2) phenotypic susceptibility profiles.

### Reagent lyophilization

The lyophilizing process involves three sequential steps: freezing, primary drying, and secondary drying. In the first step, a frozen matrix is formed, in which water converts into ice crystals and solutes are concentrated. During primary drying, we removed the frozen water by sublimation under vacuum and low-temperature conditions, followed by the removal of the remaining unfrozen water as the final step (secondary drying).

We lyophilized CRISPR assay components following previously reported strategies.^25^ For lyophilization experiments, we used a Cas13a-based *N. gonorrhoeae* detection assay targeting the *por*A gene that had previously been validated.^22^ We prepared lyophilized pellets using a modified SHINE-buffer, termed LYO buffer (50 mM HEPES pH 8·0 with 12·5% (weight per volume) sucrose and 375 mM mannitol as cryoprotectants). The master mix contains (1 x LYO buffer, 2 mM each rNTP mix, RPA pellet, 1 U/µL RNAse Inhibitor, 45 nM *Lwa*Cas13a, 1 U/µL T7 RNA Polymerase, 0·0625 µM 6U-FAM reporter (FAM-UUUUUU-quencher), 22·5 nM gRNA, 0·12 µM each RPA primer mix).

We flash-froze aliquots of the master mix in liquid nitrogen and lyophilized them overnight in a FreeZone 70040 4.5L Freeze Dryer (Labconco, United States) at −50°C and a vacuum pressure of <0.4 mbar. We stored the lyophilized pellets at −20°C for stability. We reconstituted the lyophilized pellets in resuspension buffer (60 mM Potassium Chloride (KCl), 3·5% Polyethylene Glycol 8000 (PEG-8000), 25 mM Magnesium Acetate (MgAOc), nuclease-free water).

For those experiments, we used extracted DNA from cultured isolates and performed fluorescence-based detection on the Cytation 5. We evaluated isolates in triplicates and analyzed the mean results across technical triplicates. We then evaluated the median peak fluorescence and interquartile range (IQR) as well as time to detection (1/2 peak fluorescence) and IQR across isolates comparing the lyophilized system to an aqueous control (a reaction prepared as described above).

### Data analysis

We compared mean differences in fluorescence using Student’s *t*-test among parametric data and Wilcoxon Signed-Rank Tests when data were nonparametric, with statistical significance defined as *P* < 0·05. We generated all figures using GraphPad Prism version 9·5·1 (GraphPad Software, USA).

### Ethical Considerations

The Mass General Brigham Institutional Review Board approved this study under protocols 2019P003305 and 2020P000323.

## RESULTS

### Selection and validation of primer-guide RNA pair for Cas13a-based penA assay

BADGERS produced 80 potential gRNA sequences (Supplemental Table 3). The 4 selected gRNAs had a median composite fitness score of −0.03 (IQR −0.04 to −0.02), a median composite on-target activity score of −0.70 (IQR −0.73 to −0.67) and median composite off-target activity score of −3.16 (IQR −3.37 to −2.57).

The *pen*A non-mosaic gRNA 4 paired with primer set 1 produced the highest fluorescence signal and the greatest separation between the *pen*A target and negative controls (Figure 1a). That assay correctly classified 6 *N. gonorrhoeae* isolates, including three non-mosaic and three mosaic strains (Figure 1b). Further, that assay detected non-mosaic synthetic DNA down to 3·3 copies/µL (Figure 2). We therefore selected *pen*A non-mosaic gRNA 4 and primer set 1 for further analytical validation.

**Figure 1:**
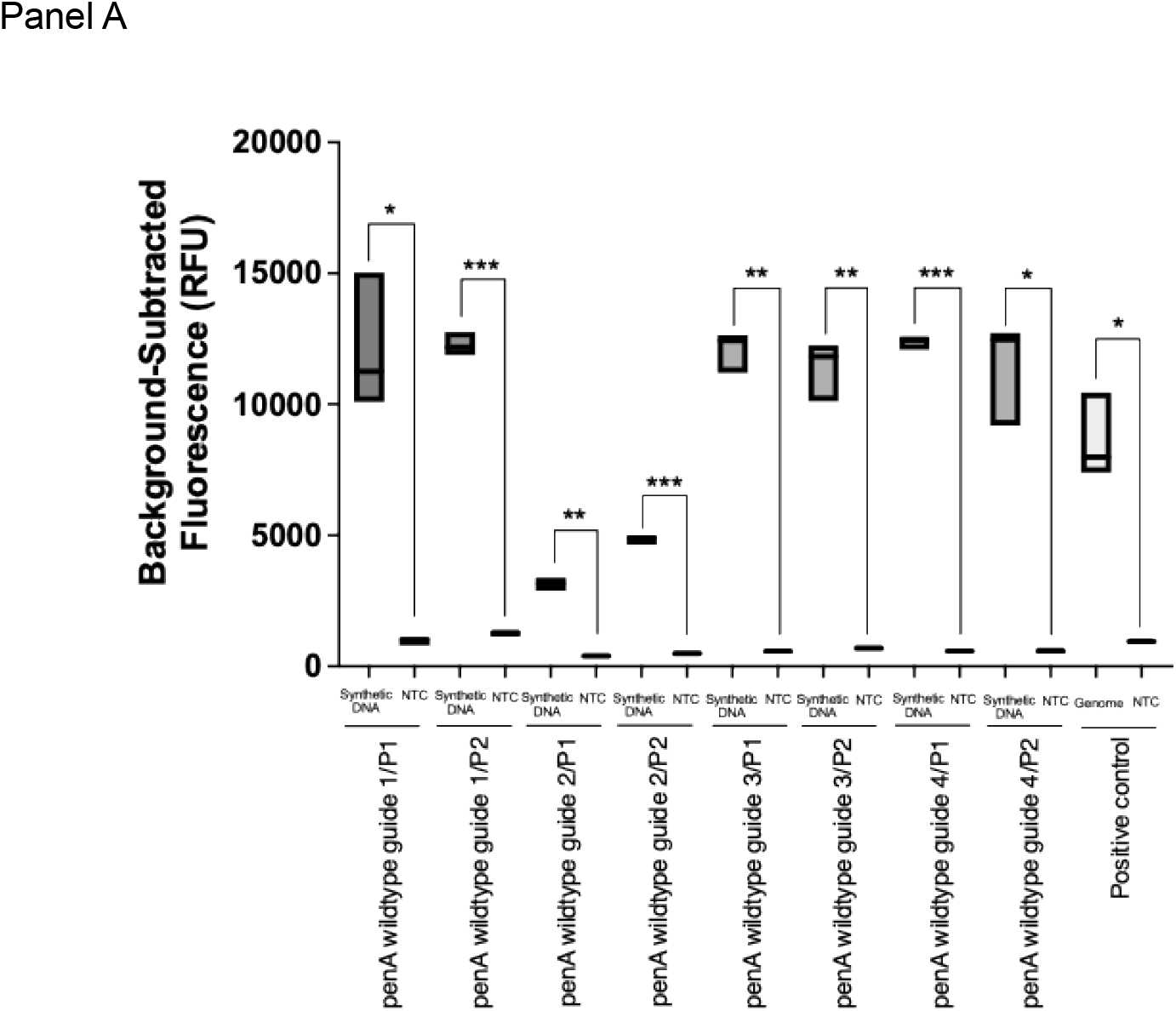
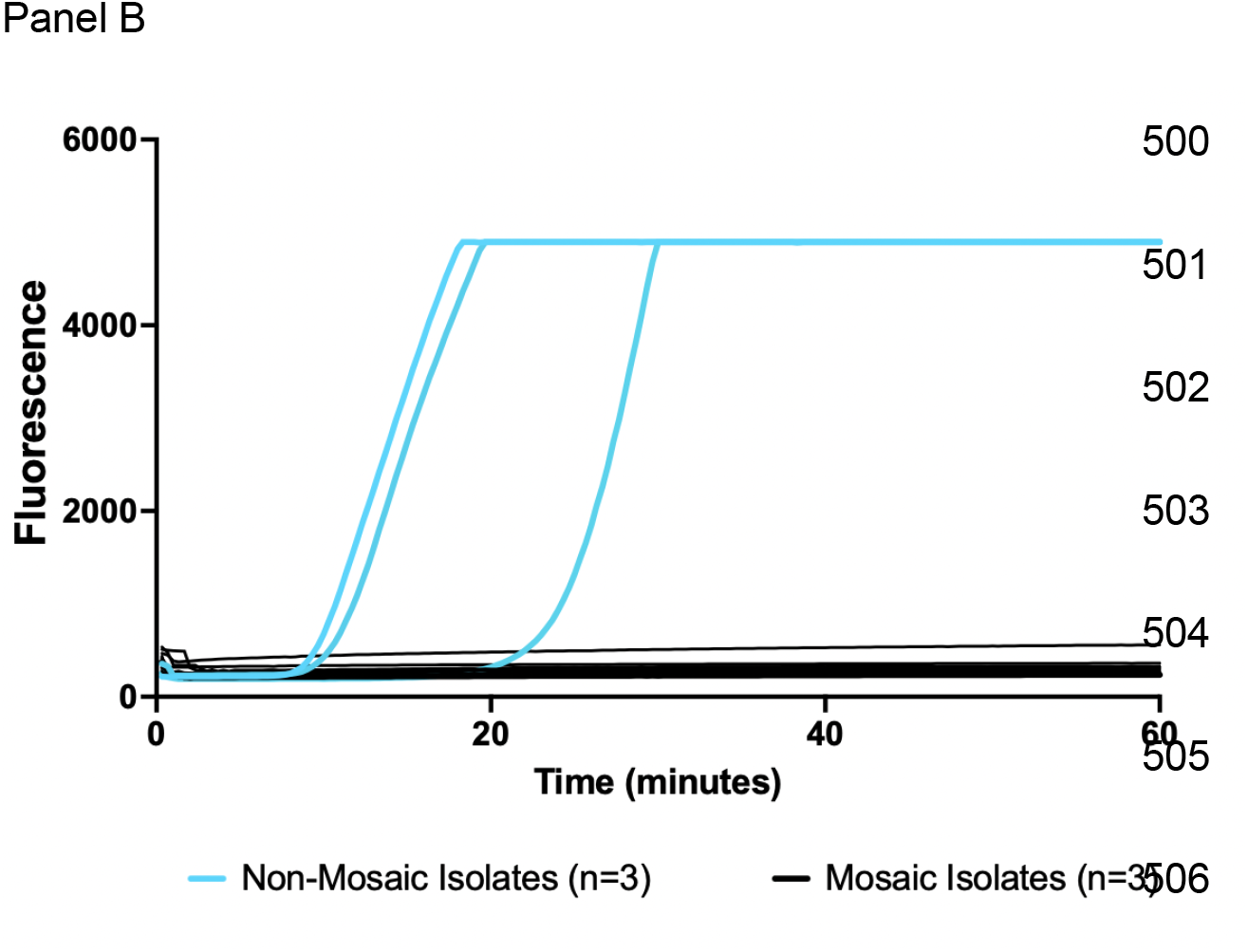
Cas13a gRNA and Primer Evaluation for Detecting Non-Mosaic *pen*A in *N. gonorrhoeae* Isolates. The figure shows the performance of four gRNAs targeting different regions of the *pen*A gene, tested on a synthetic *N. gonorrhoeae* DNA with a positive control and a negative control (NTC) using the Cytation 5 (Panel A) and Cas13a-based *pen*A mosaic assay discrimination between non-mosaic (n=3) and mosaic (n=3) *N. gonorrhoeae* isolates using the gRNA 4 - primer set 1 on the DxHub (Panel B). * p ≤ 0·05; ** p ≤ 0.01; *** p ≤ 0.001

**Figure 2:**
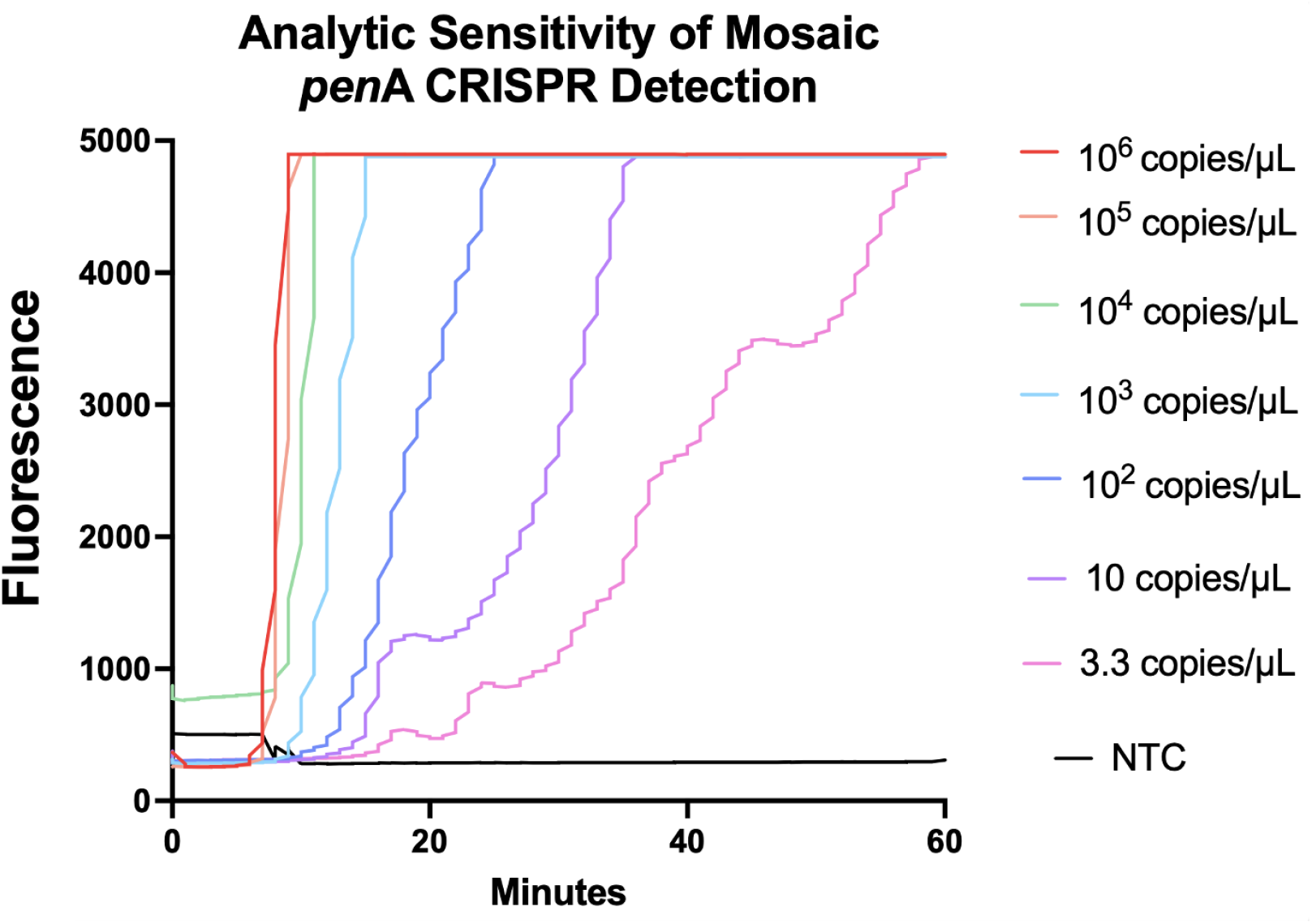
Limit of Detection of the Cas13a *pen*A Assay via Serial Dilution of Synthetic DNA. The figure shows the *in vitro* limit of detection of the Cas13a *N. gonorrhoeae pen*A assay on DxHub via ten-fold serial dilutions of a synthetic *pen*A DNA and fluorescence signals were measured in real time.

### Cas13a Determination of non-mosaic penA Genotypes Among Cultured Isolates Using the DxHub

Of the 40 isolates included, 30 had MICs ≥ 0·125 µg/mL and 10 had MICs < 0·125 µg/mL (Table). Among those isolates, the Cas13a *pen*A assay demonstrated 100% (95% CI 91.2% - 100%) concordance with qPCR. Three non-susceptible isolates (n = 2 with MIC 0·25 µg/mL; n = 1 with MIC of 0·125 µg/mL) lacked the mosaic *pen*A allele, while no susceptible isolate harbored *pen*A mosaicism. The concordance of Cas13a-based *pen*A mosaic genotyping with phenotypic susceptibility was 92·5% (37/40 isolates; 95% CI 79.6% - 98.4%). Figure 3 presents the diagnostic performance across all isolates. The median time to detection was 12 minutes (interquartile range [IQR] 5 min).

**Figure 3:**
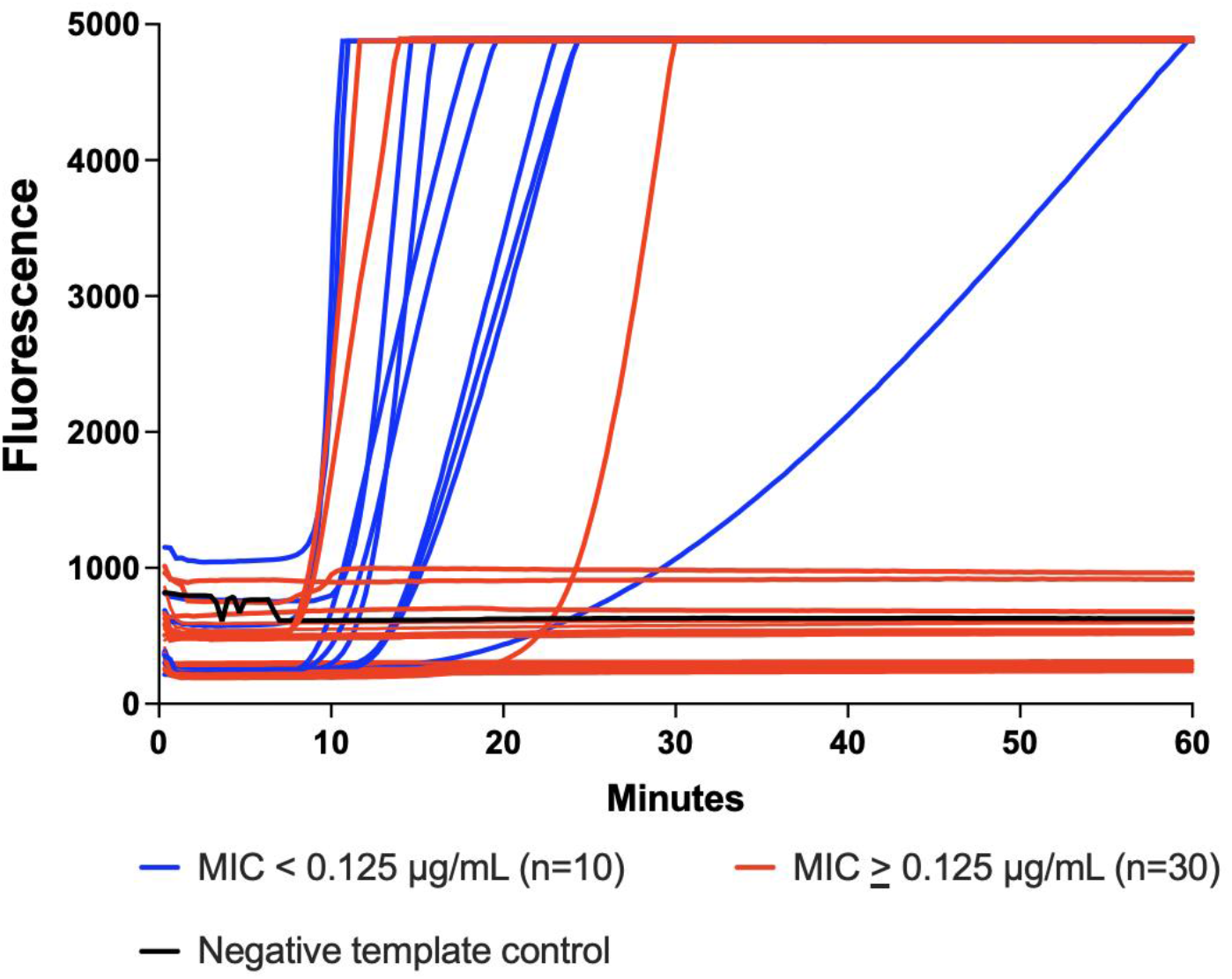
Performance of the Cas13a *pen*A Assay Among Cultured *N. gonorrhoeae* Isolates Tested on a Field-Deployable Device. The figure shows the performance of the Cas13a *pen*A assay for detecting cultured *N. gonorrhoeae* on a field-deployable platform. Blue lines represent cefixime-susceptible isolates (MIC < 0·125 µg/mL), red lines indicate cefixime non-susceptible isolates (MIC ≥ 0·125 µg/mL). Two isolates with MICs 0·25 µg/mL and one with MIC 0·125 µg/mL lacked *pen*A mosaicism. The black line represents the no-template control.

### *Cas13a-based N. gonorrhoeae* Detection Using a *Lyophilized Assay*

As a proof-of-concept, we tested a total of 12 *N. gonorrhoeae* isolates using a lyophilized Cas13a system detecting the presence of the *por*A gene. The lyophilized system detected all 12 isolates, distinguishing those from negative controls (Figure 4).The median time to detection for the lyophilized system was 45·0 minutes (IQR 40-45 minutes), while the median time to detection for the positive aqueous control was 45·0 minutes (IQR 35-55 minutes). The peak fluorescence amplitude was highest for the aqueous control (median 74,108 relative fluorescence units [RFUs]) compared with 46,802 RFUs for the lyophilized system (*p*<0.01).

**Figure 4:**
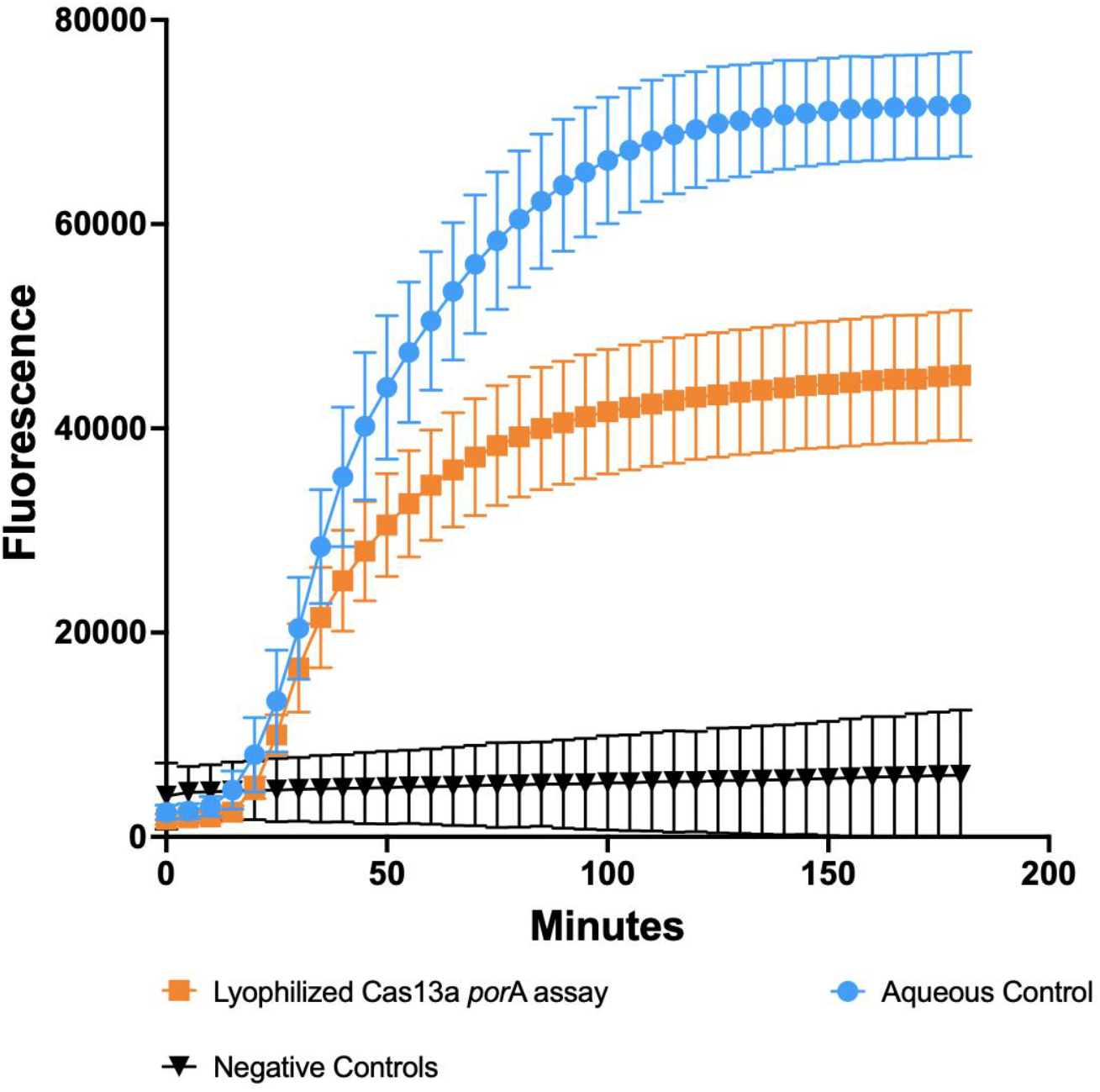
Proof-of-Concept Performance of Lyophilized Cas13a *N. gonorrhoeae* Detection Compared with Aqueous Controls Among Cultured Isolates. The figure shows the fluorescence detection on the Cytation 5 fluorometer of 12 *N. gonorrhoeae* isolates evaluated in triplicate under aqueous conditions functioning as positive controls (blue lines), lyophilized pellets containing gRNA and primers (orange lines), and negative controls (black lines). The error bars represent the 95% confidence intervals.

## DISCUSSION

This study aimed to develop a field-deployable, Cas13a-based method for rapidly determining the absence of mosaicism in the *pen*A gene of *N. gonorrhoeae*. The *pen*A mosaic assay accurately detected the absence of mosaicism compared with qPCR, which demonstrated promising correlation with phenotypic cefixime susceptibility. Integration of Cas13a assays into the DxHub may facilitate the use of such assays in field settings. Additionally, this study demonstrated promising proof-of-concept preservation of Cas13a-based *N. gonorrhoeae* detection system following lyophilization, which can permit cold-chain-independent storage and deployment of CRISPR assays in resource-limited settings.

While mosaic codons 375-377 are strongly associated with cefixime resistance, other mutations are also important predictors. Prior work developed a 6-codon algorithm (positions 375-377, 501, 542, 551) in *pen*A that predicted cefixime-decreased susceptibility in 99·5% of international isolates and 95·9% of additional strains in external datasets.^19,20^ Our results are consistent with those findings because three (out of 40) isolates demonstrated genotypic and phenotypic discordance. Another key substitution is A501V/T.^19,20^ One study recently reported the development of a CRISPR-Cas12a platform that detected both the mosaic *pen*A allele and mutations at codon 501, as well as genes for *N. gonorrhoeae* detection and a mutation associated with ceftriaxone resistance.^27^ That assay also represents a promising development towards field-deployment of genetic resistance assays. However, Cas12a necessitates separating amplification from detection for diagnostics developed for DNA organisms due to the indiscriminate trans-cleavage activity of DNA of Cas12a.^28^ Conversely, Cas13a is an RNAse, which permits one-pot amplification and detection. Future work could aim to expand the repertoire of Cas13a tests to simplify and streamline workflow.

The benefits of rapid determination of cefixime susceptibility in *N. gonorrhoeae* are numerous. First, rapid molecular resistance assay may be able to guide appropriate use and reduce the risk of treatment failures. Second, while rapid molecular assays are increasingly available for predicting ciprofloxacin resistance, incorporation of rapid susceptibility prediction to additional classes of antibiotics may have a larger impact on the delaying the emergence of ceftriaxone resistance.^18^ Further, both cefixime and ciprofloxacin are oral therapies, which are often preferable for patients over intramuscular injection. Additionally, oral antibiotics can obviate the need to return to clinic, potentially reducing losses to follow-up.^29^ Finally, prior work has demonstrated that the prevalence of *N. gonorrhoeae* infection with discordant genotypes between current sex partners is less than 3%.^30^

Our findings that lyophilized Cas13a reactions maintain functionality is also consistent with prior research.^25^ We did note a reduction in peak fluorescence and a potential slowing of reaction kinetics. Further optimization of lyophilized conditions may be able to improve the activity of the lyophilized assays; however, preservation of discriminatory capacity is a promising step towards cold-chain free CRISPR assays. Resistance-guided therapy using oral antimicrobials has the potential to thus improve patient care and facilitate expedited partner therapy – an essential component of our public health strategy to mitigate the spread of STIs. The development of low-cost, field-deployable resistance assays has the potential to extend those benefits to low-resource settings.

## LIMITATIONS

This study has several limitations. First, the sample size of *N. gonorrhoeae* isolates evaluated was small, which limits the precision of our findings. Second, while *pen*A mosaicism is an important determinant of cefixime resistance, incorporation of additional single nucleotide polymorphisms, such as those occurring at codons 501, 542, and 551, will be needed for comprehensive prediction of cefixime susceptibility. Finally, we assessed the stability and performance of lyophilized reagents under limited conditions; further studies are needed to evaluate long-term storage and robustness across a range of temperatures and real-world field conditions.

## CONCLUSION

We report on the development of a rapid, field-deployable Cas13a-based *pen*A mosaic assay that accurately detected the absence of mosaicism in the *pen*A gene of *N. gonorrhoeae*. The assay demonstrated promising correlation with phenotypic cefixime susceptibility. Integration into the portable DxHub platform could enable point-of-care use in field settings, while proof-of-concept lyophilization could support cold-chain-independent deployment, offering a practical tool for resistance-guided therapy and antimicrobial stewardship, particularly in resource-limited settings.

## Funding and Acknowledgements

This work was supported by the National Institutes of Health’s National Institute of Allergy and Infectious Diseases (K23AI182453 to LAB), the Brigham Research Institute, and the Hearst Family Foundation.

---

**Table :** Phenotypic cefixime susceptibility profile of 40 *N. gonorrhoeae* isolates evaluated for *pen*A mosaicism using qPCR and a Cas13a-based Assay.

| ID | Cefixime MIC (µg/mL)<sup>a</sup> | Resistance interpretation | *penA* by qPCR | *penA* by Cas13a<sup>†</sup> | Cas13a Concordance |  |
| --- | --- | --- | --- | --- | --- | --- |
|  |  |  |  |  | PCR | Phenotype |
| CCC001 | 0.25 | Resistant | Non-mosaic | Non-mosaic | Yes | No |
| CCC018 | 0.25 | Resistant | Non-mosaic | Non-mosaic | Yes | No |
| FQ009 | 0.125 | Intermediate | Non-mosaic | Non-mosaic | Yes | No |
| FQ023 | ≤0.015 | Susceptible | Non-mosaic | Non-mosaic | Yes | Yes |
| FQ027 | 0.06 | Susceptible | Non-mosaic | Non-mosaic | Yes | Yes |
| FQ030 | ≤0.015 | Susceptible | Non-mosaic | Non-mosaic | Yes | Yes |
| FQ038 | ≤0.015 | Susceptible | Non-mosaic | Non-mosaic | Yes | Yes |
| FQ043 | 0.06 | Susceptible | Non-mosaic | Non-mosaic | Yes | Yes |
| FQ047 | ≤0.015 | Susceptible | Non-mosaic | Non-mosaic | Yes | Yes |
| FQ049 | 0.06 | Susceptible | Non-mosaic | Non-mosaic | Yes | Yes |
| FQ056 | 0.06 | Susceptible | Non-mosaic | Non-mosaic | Yes | Yes |
| FQ059 | ≤0.015 | Susceptible | Non-mosaic | Non-mosaic | Yes | Yes |
| FQ093 | 0.06 | Susceptible | Non-mosaic | Non-mosaic | Yes | Yes |
| CCC006 | 0.25 | Resistant | Mosaic | Mosaic | Yes | Yes |
| CCC009 | 2 | Resistant | Mosaic | Mosaic | Yes | Yes |
| DDD004 | 2 | Resistant | Mosaic | Mosaic | Yes | Yes |
| DDD021 | 0.5 | Resistant | Mosaic | Mosaic | Yes | Yes |
| DDD022 | 1 | Resistant | Mosaic | Mosaic | Yes | Yes |
| DDD035 | 0.5 | Resistant | Mosaic | Mosaic | Yes | Yes |
| DDD044 | 1 | Resistant | Mosaic | Mosaic | Yes | Yes |
| DDD050 | 0.5 | Resistant | Mosaic | Mosaic | Yes | Yes |
| EEE004 | 2 | Resistant | Mosaic | Mosaic | Yes | Yes |
| FQ091 | 0.125 | Intermediate | Mosaic | Mosaic | Yes | Yes |
| FQ041 | 0·25 | Resistant | Mosaic | Mosaic | Yes | Yes |
| FQ045 | 0·25 | Resistant | Mosaic | Mosaic | Yes | Yes |
| FQ048 | 0·25 | Resistant | Mosaic | Mosaic | Yes | Yes |
| FQ050 | 0·25 | Resistant | Mosaic | Mosaic | Yes | Yes |
| FQ053 | 0·25 | Resistant | Mosaic | Mosaic | Yes | Yes |
| FQ069 | 0·25 | Resistant | Mosaic | Mosaic | Yes | Yes |
| FQ071 | 0·25 | Resistant | Mosaic | Mosaic | Yes | Yes |
| FQ057 | 0·25 | Resistant | Mosaic | Mosaic | Yes | Yes |
| FQ060 | 0·25 | Resistant | Mosaic | Mosaic | Yes | Yes |
| FQ062 | 0·25 | Resistant | Mosaic | Mosaic | Yes | Yes |
| FQ074 | 0·25 | Resistant | Mosaic | Mosaic | Yes | Yes |
| FQ076 | 0·25 | Resistant | Mosaic | Mosaic | Yes | Yes |
| FQ078 | 0·25 | Resistant | Mosaic | Mosaic | Yes | Yes |
| FQ081 | 0·25 | Resistant | Mosaic | Mosaic | Yes | Yes |
| FQ083 | 0·25 | Resistant | Mosaic | Mosaic | Yes | Yes |
| FQ085 | 0·25 | Resistant | Mosaic | Mosaic | Yes | Yes |
| FQ088 | 0·25 | Resistant | Mosaic | Mosaic | Yes | Yes |

<sup>a</sup> Minimum inhibitory concentration
† Cas13a detection indicates the presence of a non-mosaic *penA* allele

